# Cross-linking/Mass Spectrometry Combined with Ion Mobility on a timsTOF Pro Instrument for Structural Proteomics

**DOI:** 10.1101/2021.03.26.437136

**Authors:** Christian H. Ihling, Lolita Piersimoni, Marc Kipping, Andrea Sinz

**Affiliations:** Department of Pharmaceutical Chemistry & Bioanalytics, Institute of Pharmacy, Kurt-Mothes-Str. 3, D-06120 Halle (Saale), Germany; Center for Structural Mass Spectrometry, Kurt-Mothes-Str. 3, D-06120 Halle (Saale), Germany

**Keywords:** Chemical Cross-Linking, Ion Mobility, Mass Spectrometry, Protein 3D-Structure, Protein Interaction Networks

## Abstract

The combination of cross-linking/mass spectrometry (XL-MS) and ion mobility is still underexplored for conducting protein conformational and protein-protein interaction studies. We present a method for analyzing cross-linking mixtures on a timsTOF Pro mass spectrometer that allows separating ions based on their gas phase mobilities. Cross-linking was performed with three urea-based MS-cleavable cross-linkers that deliver distinct fragmentation patterns for cross-linked species upon collisional activation. The discrimination of cross-linked species from non-cross-linked peptides was readily performed based on their collisional cross sections. We demonstrate the general feasibility of our combined XL-MS/ion mobility approach for three protein systems of increasing complexity: (i) Bovine serum albumin, (ii) *E. coli* ribosome, and (iii) HEK293T cell nuclear lysates. We identified a total of 508 unique cross-linking sites for BSA, 868 for the *E. coli* ribosome, and 1,623 unique cross-links for nuclear lysates, corresponding to 1,088 intra- and 535 interprotein interactions and yielding 564 distinct protein-protein interactions. Our results underline the strength of combining XL-MS with ion mobility not only for deriving 3D-structures of single proteins, but also for performing system-wide protein interaction studies.

**TOC Graphic:** 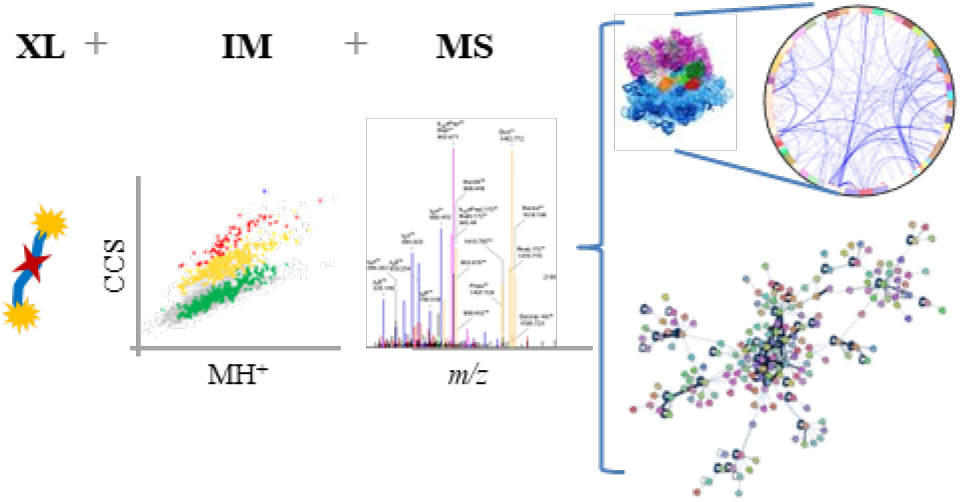

## Introduction

Investigating the three-dimensional structures of proteins is usually performed by NMR spectroscopy, X-ray crystallography, and cryo-electron microscopy (EM). In recent years, integrated methods of structural biology have come into focus as they combine different low-resolution techniques, such as small-angle X-ray scattering (SAXS) or cross-linking/mass spectrometry (XL-MS) with partial high-resolution structural data and computational modeling. The XL-MS technique has proven extremely useful in combination with cryo-EM to resolve the structures of large protein assemblies, so-called “molecular machines”^1^.

XL-MS has evolved in the last 20 years into an alternative strategy to structurally resolve protein conformations and protein-protein interactions^2, 3, 4, 5, 6, 7, 8, 9, 10^. It allows deriving a set of structurally defined interactions, by covalently connecting pairs of functional groups within a protein. From the distance constraints imposed by the chemical cross-links, distance maps can be generated, which will serve as basis for inferring 3D-structural information of proteins and protein assemblies. The outstanding properties of XL-MS include the minimal sample requirements and the theoretically unlimited size of the protein assemblies that are to be studied. The chemical XL-MS approach has received growing interest among system biologists as the technique is increasingly being applied to map protein interaction networks in cells, organs, and tissue^11, 12, 13, 14^.

Combining XL-MS with ion mobility mass spectrometry (IM-MS) is a still a largely underexplored field. In IM-MS ions are separated based on their collisional cross sections (CCS, Ω), which depend on size and shape, and charge states of the molecules. Recently, a commercially available instrument combining trapped ion mobility spectrometry (TIMS) with time-of-flight (TOF) mass analysis (timsTOF Pro, Bruker Daltonik) using PASEF (parallel accumulation serial fragmentation) allowed the identification of more than 6,000 proteins during LC/MS analyses^15, 16, 17^. An optimized data analysis workflow for timsTOF Pro data using the software tools MSFragger and IonQuant has been reported, improving protein identification, sensitivity, quantification accuracy, and runtimes in proteomics experiments^18^. The timsTOF Pro mass spectrometer has been employed recently in a XL-MS study by the groups of Heck and Scheltema using the enrichable PhoX cross-linker^19^. In that study, an immobilized metal ion chromatography (IMAC) enrichment of cross-linker-containing peptides was performed before the MS analysis. The various crossed species generated in a cross-linking experiment comprise the following three types: (i) intrapeptide cross-links (amino acids within one peptide are cross-linked), (ii) interpeptide cross-links (amino acids between two peptides are cross-linked), and (iii) peptides that are modified by a partially hydrolyzed cross-linker, which are commonly referred to as “dead-end” cross-links or “mono-links”. The major aim of that previous study^19^ was to separate inter- and intrapeptide cross-links on one side from “dead-end” products on the other side as the latter ones do not yield direct distance information on interaction sites in proteins. The selection of cross-linked peptides was performed based on their characteristic CCS by applying a strategy termed “collisional cross-section assisted precursor selection (caps-PASEF)”. In caps-PASEF, a specific region, i.e. a polygon, is defined in the distribution of monoisotopic mass versus CCS, including specific CCS values that distinguish intra- and interpeptide cross-linked species from “dead-end” products, so only “true” cross-linked peptides are selected for fragmentation.

In our study, we applied a different approach relying on urea-based MS-cleavable cross-linking principles. This class of cross-linkers originates from the DSBU (disuccinimidyl dibutyric urea) reagent that has been developed in our group_20_. DSBU has been successfully used in the past for conformational studies of proteins as well as for mapping system-wide protein interaction networks^3, 6, 14, 20^. Here, we employ the three cross-linkers DSBU, DSPU (disuccinimidyl dipropionic urea), and DSAU (disuccinimidyl diacetic urea)^21^. All three cross-linkers are amine-reactive *N*-hydroxysuccinimide (NHS) esters and contain a central, MS-labile urea group with C4 to C2 spacers at each side of the urea moiety (Scheme 1). NHS esters mainly react with lysine residues, but are also reactive towards serine, threonine, and tyrosine residues in proteins^3^. In this study, we investigate the benefits of IM-MS to efficiently analyze *<u>all</u>* cross-linked species (intra- and interpeptide as well as “dead-end” cross-links), while non-modified, linear tryptic peptides are still present in the sample mixture. To cover a broad range of protein systems, we chose a single 66 kDa-protein (BSA), a large protein assembly (*E. coli* ribosome), and a complex protein mixture (HEK293T cell nuclear lysates) to develop a generally applicable strategy combining IM-MS and XL-MS for an efficient and fast analysis of cross-linked products.

## EXPERIMENTAL

### Reagents

All chemicals and solvents were purchased from Sigma-Aldrich, Roth, and VWR. Cell culture reagents were from Gibco. The MS-cleavable cross-linkers DSBU (disuccinimidyl dibutyric urea, cat. no. 1881355), DSPU (disuccinimidyl dipropionic urea, cat. no. 1881356), and DSAU (disuccinimidyl diacetic urea, cat. no. 1881357) were provided by Bruker Daltonik. Each cross-linker (10 mg supplied in 5 x 2 mg vials) was aliquoted under argon.

### BSA and Ribosome Cross-linking

All cross-linking reaction mixtures were analyzed in three (BSA) or four replicates (ribosome). BSA was purchased from Sigma-Aldrich (cat. no. A7638). A 10 μM BSA solution in 50 mM aqueous HEPES (2-[4-(2-hydroxyethyl)-piperazine-1-yl]ethanesulfonic acid), pH 7.5, was cross-linked at room temperature (RT) for 1 hr with DSBU, DSPU, and DSAU (each at 25, 50, and 125-fold molar excess) in separate samples. All cross-linkers were dissolved in neat DMSO at a concentration 25 mM immediately before adding them to the BSA solution. *E. coli* ribosome was purchased from New England Biolabs (NEB, cat. no. P0763S20) and rebuffered in 20 mM HEPES, 10 mM Mg(Ac)_2_, 30 mM KCl, pH 7.6, using Zeba spin columns (Thermo Fisher Scientific). The concentration was adjusted to 1.25 mg/mL total protein. For the cross-linking reactions, DSBU was dissolved in DMSO (25 mM stock solution) and added to a final concentration of 1 mM to the *E. coli* ribosome (according to a molar ratio of 2:1 between DSBU and lysine residues). The reaction was allowed to proceed for 1 hr at RT.

### Cell Culture, Isolation of Nuclei, and Cross-linking

HEK293T cells (ATCC) were cultured in VDMEM medium supplemented with 10% (v/v) fetal bovine serum (Gibco). Three 10-cm plates of 65/70% confluent cells were used as replicates. Cells were washed once with phosphate buffered saline (PBS), scraped in PBS, and then counted. From each sample, aliquots containing ~2 million cells were collected and centrifuged at 1,500 rpm for 3 min. The samples were washed again in 2 ml of PBS, transferred in Eppendorf tubes and centrifuged at 4,000 rpm for 3 min. The nuclear pellets were resuspended in 200 μl of lysis buffer (10 mM HEPES, 150 mM KCl, 5 mM MgCl_2_, 80 μg/ml digitonin, pH 7.5), incubated at RT for 7 min and centrifuged as above. The nuclear pellets were washed twice with 200 μl of lysis buffer and finally resuspended in 100 μl of cross-linking buffer (10 mM HEPES, 150 mM KCl; 5 mM MgCl_2_, pH 7.7). DSBU was resuspended in DMSO, added to a final concentration of 0.5 mM. The mixture was incubated at RT for 10 min. The cross-linking reaction was quenched with a 20 mM final concentration of ammonium bicarbonate. In order to break the nuclei, sodium dodecyl sulfate (SDS) was added to a final concentration of 1% (w/v) together with 12.5 units of benzonase (Merck); after 5 min of incubation at RT, samples were centrifuged for 2 min at 15,000 *x g* at 4°C. The supernatants containing the nuclear lysates were used for further analysis.

### Enzymatic Digestion

All cross-linked BSA samples and nuclear lysates were dried down to a volume of 3-5 μl in a SpeedVac concentrator. 25 μl of 8 M urea in 400 mM ammonium bicarbonate was added to each sample. After 5 min of sonication, samples were incubated with DTT (6.4 mM) for 30 min at 56°C and iodoacetamide (12.5 mM) for 20 min at RT in the dark. Alkylation was quenched by adding 9 mM of DTT. After diluting the BSA samples to 1 M urea with water, they were incubated at 37°C and digested with trypsin (Promega) overnight. For proteolytic digestion of nuclear lysates, samples were preincubated at 37°C for 4 hours with 2 μg AspN (NEB) before tryptic digestion was performed at 37°C overnight. 50 μl of cross-linked *E. coli* ribosome was proteolytically digested with the SMART Digest Trypsin Kit (Thermo Fisher Scientific). 150 μL of SMART Digest buffer were added and the resulting solutions were incubated at 70°C for 3 hrs. After cooling down, the samples were centrifuged and supernatants were collected. Peptides were further incubated with DTT (4 mM) at 56°C for 30 min and iodoacetamide (8 mM) at RT for 20 min in the dark. Alkylation was quenched by adding 4 mM of DTT. All proteolytic digestions were stopped by adding TFA.

### Size Exclusion Chromatography of Nuclear Lysates

All three replicates of digested nuclear lysates were subjected to size exclusion chromatography (SEC) on an ÄKTA FPLC system (equipped with UPC-900, P-920 pump, and Frac-950 fraction collector, Amersham Biosciences) for prefractionation before LC/MS/MS analysis was performed^14^. Peptides were separated on a Superdex peptide 3.2/300 column (GE Healthcare) using a 2% (v/v) ACN, 0.1 % (v/v) TFA buffer at flow rate of 50 μl/min. 2-min fractions were collected over one hour; fractions B1 to B7 were selected for subsequent LC/MS/MS analyses.

### LC/MS/MS

Digested samples of BSA and the *E. coli* ribosome were analyzed in three or four technical replicates, nuclear lysate samples in three biological replicates by LC/MS/MS on an UltiMate 3000 RSLC nano-HPLC system (Thermo Fisher Scientific) that was coupled to a timsTOF Pro mass spectrometer equipped with CaptiveSpray source (Bruker Daltonics). Peptides were trapped on a C18 column (precolumn Acclaim PepMap 100, 300 μm × 5 mm, 5μm, 100 Å (Thermo Fisher Scientific) and separated on a μPAC 50 column (PharmaFluidics). After trapping, peptides were eluted by a linear 90-min (for BSA and ribosome) or 240-min (for nuclear lysates) water-ACN gradient from 3% (v/v) to 35% (v/v) ACN. For elution, a flow gradient was employed ranging from 900 nl/min to 600 nl/min in 15 min, followed by a constant flow rate of 600 nl/min. The column was washed at a flow rate of 600 nl/min with the following gradient: 35% (v/v) to 85% (v/v) ACN (5 min), 85% (v/v) ACN (5 min), 85% (v/v) to 3% (v/v) ACN (5 min). All separations were performed at RT.

For the timsTOF Pro settings, the following parameters were adapted, starting from the PASEF method for standard proteomics. The values for mobility-dependent collision energy ramping were set to 95 eV at an inversed reduced mobility (1/k_0_) of 1.6 Vs/cm^2^ and 23 eV at 0.73 Vs/cm^2^. Collision energies were linearly interpolated between these two 1/k_0_ values and kept constant above or below. No merging of TIMS scans was performed. Target intensity per individual PASEF precursor was set to 20,000. The scan range was set between 0.6 and 1.6 Vs/cm^2^ with a ramp time of 166 ms. 14 PASEF MS/MS scans were triggered per cycle (2.57 s) with a maximum of 7 precursors per mobilogram. Precursor ions in an *m/z* range between 100 and 1700 with charge states ≥ 3+ and ≤ 8+ were selected for fragmentation. Active exclusion was enabled for 0.4 min (mass width 0.015 Th, 1/k_0_ width 0.015 Vs/cm^2^).

### Data Analysis

Identification of cross-links was performed with MeroX 2.0.1.4^6^. The following settings were applied: Proteolytic cleavage: *C*-terminal at Lys and Arg for BSA and ribosome (3 missed cleavages were allowed), for nuclear lysates also *N-*terminal at Asp and Glu (3 missed cleavages were allowed); peptide lengths of 5 to 50 amino acids; modifications: alkylation of Cys by iodoacetamide (fixed), oxidation of Met (variable); cross-linker specificity: Lys, Ser, Thr, Tyr, *N*-terminus (sites 1 and 2 for BSA and ribosome); Lys and *N*-terminus and Lys, Ser, *N*-terminus (sites 1 and 2, respectively, for nuclear lysates); search algorithm: RISEUP mode with up to two missing ions (for BSA and ribosome) or proteome-wide mode with minimum peptide score of 5 (for nuclear lysates); precursor mass accuracy: 15 ppm; fragment ion mass accuracy: 25 ppm; signal-to-noise ratio > 2 (for BSA and ribosome) or >1.5 (for nuclear lysates); precursor mass correction enabled; 10% intensity as prescore cut-off; 1% false discovery rate (FDR) cut-off, and minimum score cut-off: 20. Proteome Discoverer 2.4 was used for the identification of proteins present in nuclear lysates (all relevant fractions from the three biological replicates were combined) using the human proteome database (uniprot.org; accession date: Nov. 15, 2020; 20,315 entries). All protein IDs with FDR < 5% were combined in a sub-database (2803 entries: 2654 entries at 1% FDR, 149 entries at 5% FDR) for subsequent cross-link searches with MeroX. MS data have been deposited to the ProteomeXchange Consortium via the PRIDE partner repository with the project accession PXD025023, username, password: v6B22Auu.

### Visualization of Cross-links

Cross-links in *E. coli* ribosome were visualized in xVis^22^ and selected proteins from HEK293T nuclear lysates were visualized in PyMOL (The PyMOL Molecular Graphics System, Version 2.0 Schrödinger, LLC) and xiNET^23^. Protein-protein interaction networks in nuclear lysates were visualized with Cytoscape 3.8.2^24^.

## RESULTS and DISCUSSION

### Ion Mobility Mass Spectrometry (IM-MS) for Cross-link Analysis

The combination of XL-MS and IM-MS presents an appealing option for improving the identification of cross-links in structural proteomics studies based on their CCS. The most useful cross-link species for deriving structural information on protein 3D-structures and for mapping protein-protein interactions networks are interpeptide cross-links. However, these are usually low-abundant species in cross-linking reaction mixtures. Interpeptide cross-links are accompanied by intrapeptide cross-links and peptides that are modified by a partially hydrolyzed cross-linker – also referred to as “dead-end” cross-links or “mono-links”. The latter species do not provide direct distance information, but yield important insights into the overall topology of proteins based on solvent accessibility of specific amino acid side chains. As such, combining the information of all three cross-linked species, including “dead-end” cross-links, is highly valuable as input for a subsequent computational modeling of protein 3D-structures.

The greatest challenge in XL-MS studies is the discrimination of non-reacted, linear peptides from cross-linked species. IM-MS exhibits a distinct advantage here as cross-linked species are easily distinguished from linear tryptic peptides based on their CCS. As shown in the distributions of monoisotopic mass versus CCS for digested *E. coli* ribosome and nuclear HEK293T cell lysate, cross-linked peptides show a clearly distinct IM behavior compared to non-modified peptides (Figure 1). Interestingly, a number of cross-linked species show several different CCS values (Figure 1B), indicating that cross-linked peptides can adopt various conformations in the gas phase. Another aspect is that the specifically cross-linked amino acids may differ between isomeric cross-linking products, resulting in different CCS.

**Figure 1:**
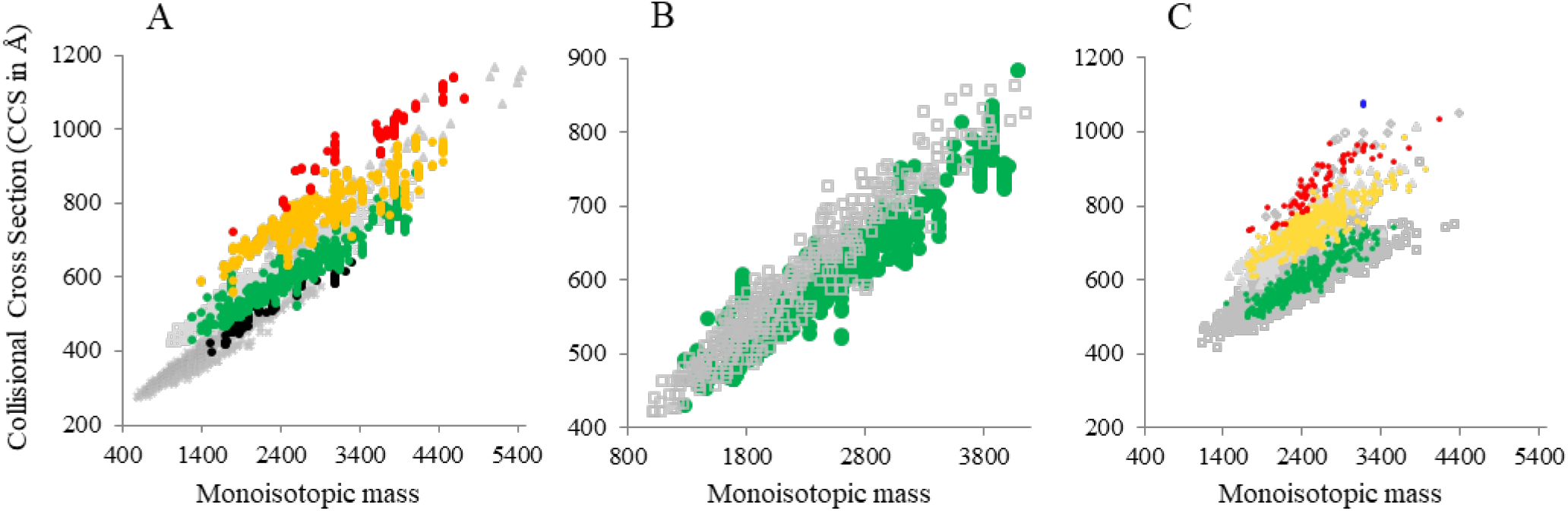
Distributions of monoisotopic mass versus CCS. (A) Cross-linked and digested *E. coli* ribosome, (B) zoom-in view on 3+ charge states of cross-linked and linear peptides from *E. coli* ribosome, and (C) digested HEK293T cell nuclear lysate. Cross-linked species (intra-, inter-peptide, and “dead-end” cross-links) are color-coded by their charge states (• 2+, • 3+, • 4+, • 5+, • 6+); linear peptides are shown in grey (× 2+, ⧠ 3+, Δ 4+, ◊ 5+, ○ 6+).

In our analyses, ions with charge states of 1+ and 2+ were excluded from fragmentation, while ions with charge states between 3+ to 8+ were considered as potential cross-linked products. When taking a closer look at the CCS of 3+ charge states of linear tryptic peptides versus cross-linked peptides, a clear separation is visible in the distributions of monoisotopic mass versus CCS (Figure 1B). This would allow an extraction of cross-linked species with identical charge states based on their CCS.

One goal of this study was to define general CCS characteristics of all cross-linked species that will later allow applying general selection procedures to separate cross-linked from non-cross-linked species. In the caps-PASEF approach described recently^19^, only intra- and interpeptide cross-links were selected for fragmentation, while “dead-end” cross-links were excluded. A specific region (polygon) of CCS values was defined in the distributions of monoisotopic mass versus CCS, in which the ions mainly correspond to cross-linked peptides that were then fragmented. From the CCS distributions of cross-linked ribosome and nuclear lysate samples (Figures 1A and 1C) it is clearly visible that the CCS values of cross-linked species follow a similar trend for all protein systems studied herein. This would theoretically allow assigning specific CCS regions for caps-PASEF. However, defining a single polygon for CCS values of cross-linked species (intra- and interpeptide as well as “dead-end” cross-links) would not be broadly applicable for single proteins, protein assemblies and system-wide protein-protein networks as cross-linked and linear peptides with identical charge states overlap in their CCS. Therefore, different polygons would have to be specified for each sample type. As stated before, our aim was to also include also “dead-end” cross-links in our analyses as these are providing valuable structural information on protein topology and should not be dismissed. Therefore, we chose not to define a polygon in the distributions of monoisotopic mass versus CCS for selecting precursor ions with specific CCS values (caps-PASEF) as the speed of the timsTOF Pro instrument allows a fast analysis of all ions, without losing potentially cross-linked species.

The main improvements regarding the analytical depth obtained in our experiments with the timsTOF Pro mass spectrometer are related to sensitivity and speed enhancement due to the IM option. These developments are impressive. In fact, we found the numbers of cross-linked products to be higher than those obtained with a Q-Exactive Plus instrument, which has been our instrument of choice for XL-MS analyses during the last years. On average, 298 unique cross-linking sites were identified for BSA (cross-linked with a 50-fold molar excess of DSBU) on a timsTOF Pro instrument (Table 1), compared to an average number of 198 unique cross-linking sites identified on a Q-Exactive Plus mass spectrometer.

**Table 1:**
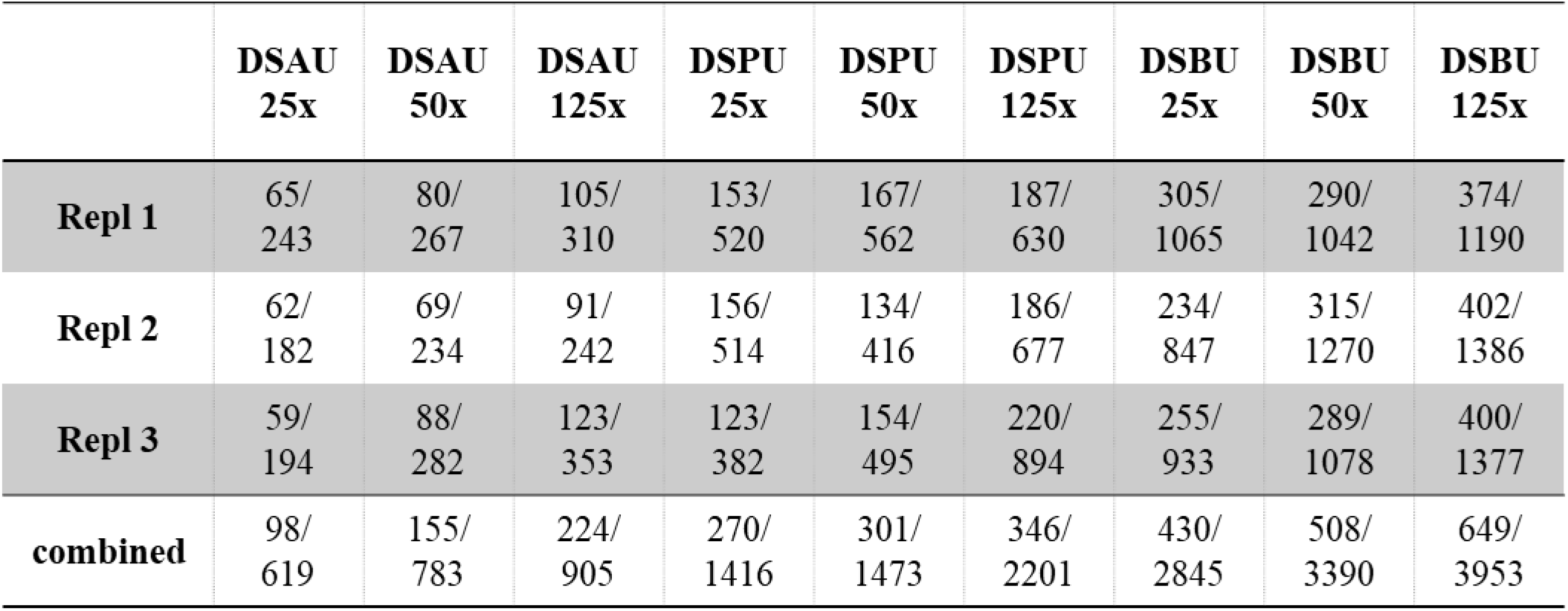
Unique cross-linking sites and cross-link spectral matches (CSMs) for BSA. Numbers are given for three replicates for all three cross-linkers and three concentrations as well as the combined numbers; unique cross-linking site/CSMs.

|  | **DSAU**<br>**25x** | **DSAU**<br>**50x** | **DSAU**<br>**125x** | **DSPU**<br>**25x** | **DSPU**<br>**50x** | **DSPU**<br>**125x** | **DSBU**<br>**25x** | **DSBU**<br>**50x** | **DSBU**<br>**125x** |
| --- | --- | --- | --- | --- | --- | --- | --- | --- | --- |
| **Repl 1** | 65/<br>243 | 80/<br>267 | 105/<br>310 | 153/<br>520 | 167/<br>562 | 187/<br>630 | 305/<br>1065 | 290/<br>1042 | 374/<br>1190 |
| **Repl 2** | 62/<br>182 | 69/<br>234 | 91/<br>242 | 156/<br>514 | 134/<br>416 | 186/<br>677 | 234/<br>847 | 315/<br>1270 | 402/<br>1386 |
| **Repl 3** | 59/<br>194 | 88/<br>282 | 123/<br>353 | 123/<br>382 | 154/<br>495 | 220/<br>894 | 255/<br>933 | 289/<br>1078 | 400/<br>1377 |
| **combined** | 98/<br>619 | 155/<br>783 | 224/<br>905 | 270/<br>1416 | 301/<br>1473 | 346/<br>2201 | 430/<br>2845 | 508/<br>3390 | 649/<br>3953 |

### Cross-links in Bovine Serum Albumin

For cross-linking BSA, all three MS-cleavable urea-based cross-linkers DSBU, DSPU, and DSAU with spacer lengths between 12.5 and 7.7 Å (Scheme 1) were employed, thereby mapping different distances in BSA. We analyzed each cross-linking sample in triplicate to get insights into the technical reproducibility of our approach. In total, 430, 508, and 649 unique cross-linking sites were identified with DSBU (25-, 50-, and 125-fold molar excess), 270, 301, and 346 unique cross-linking sites were identified with DSPU (25-, 50-, and 125-fold molar excess), and 98, 155, and 224 unique cross-linking sites were identified with DSAU (25-, 50-, and 125-fold molar excess; Table 1). Monomeric and dimeric forms of BSA coexist in all sources of commercially available BSA, including the one used in this study^3^. Therefore, cross-links identified upon *in-solution* digestion might derive from BSA monomers and dimers.

Analysis of BSA cross-linked peptides with the MeroX software indicates the efficient fragmentation of cross-linked peptides by the timsTOF Pro instrument. It should be noted that the MeroX software allowed assigning fragment ion mass spectra acquired by the timsTOF Pro mass spectrometer without any modifications in the software although MeroX had originally been developed and optimized for orbitrap instruments (Figure 2).

**Figure 2:**
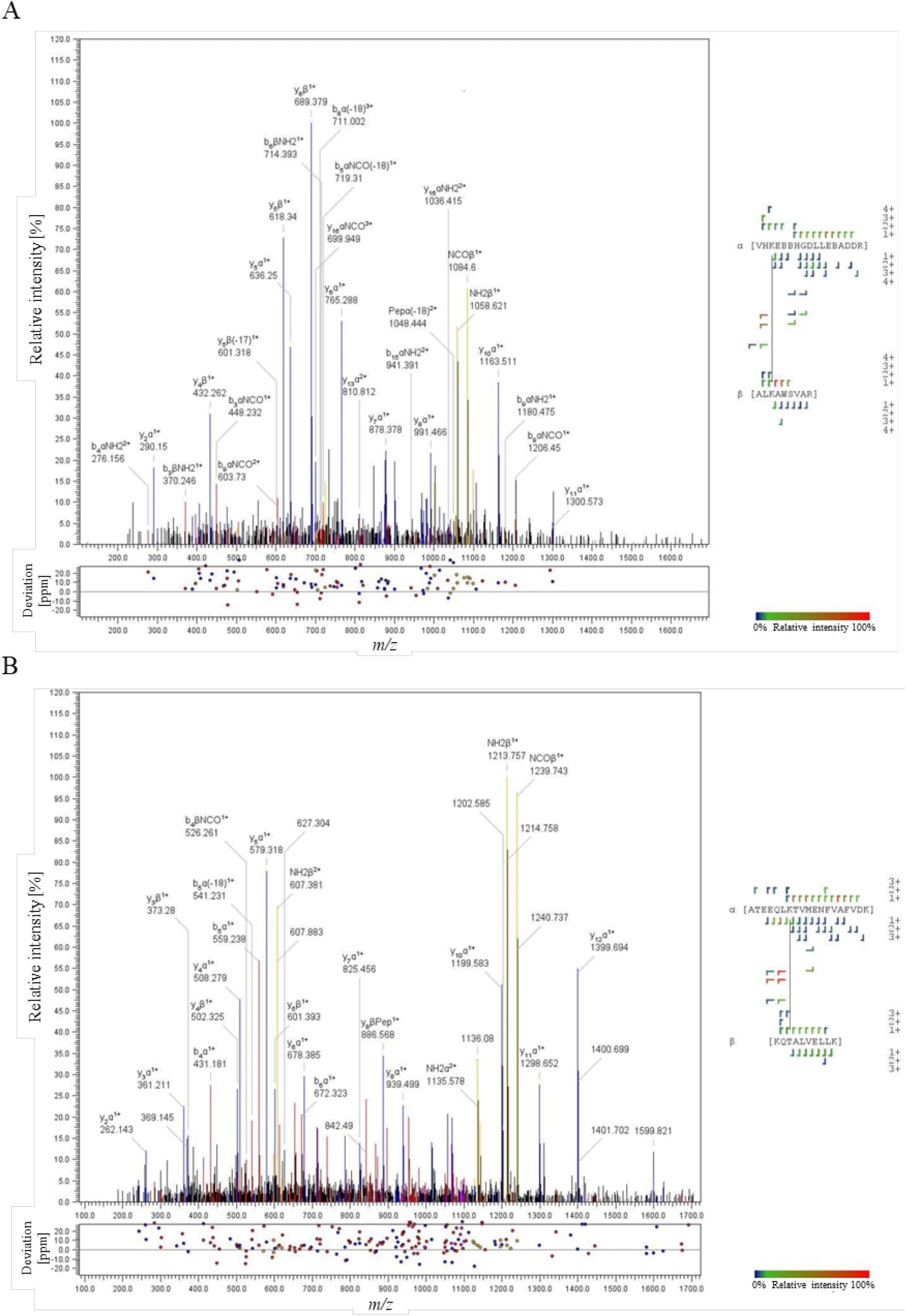
Assignment of BSA cross-linked peptides with the MeroX software. Cross-linking was performed with 50-fold molar excess of (A) DBAU and (B) DSPU. The characteristic fragment ions of the DBSU cross-linker are shown in the fragment ion mass spectra in yellow, b- and y-type ions are presented in blue and red, respectively.

The overall overlap of cross-links in BSA between the three different cross-linkers DSBU, DSPU, and DSAU is presented in Figure 3. A comparison of all cross-linking experiments performed (three technical replicates, three cross-linkers, each cross-linker used at three different concentrations) is presented in SI Figure S1. DSBU with a spacer length of 12.5 Å yields the highest number of unique cross-linking sites. Strikingly, for the shortest cross-linker applied herein, DSAU, we found the lowest number of cross-links, which is probably caused in part by a base-catalyzed intramolecular cyclization of the urea moiety (personal communication Vaclav Matousek). The overlap of unique cross-linking sites differs between 15.2% and 31.6% among single runs (SI Figure S1), while the overlap of unique cross-linking sites in all experiments combined is ~10% (Figure 3).

**Figure 3:**
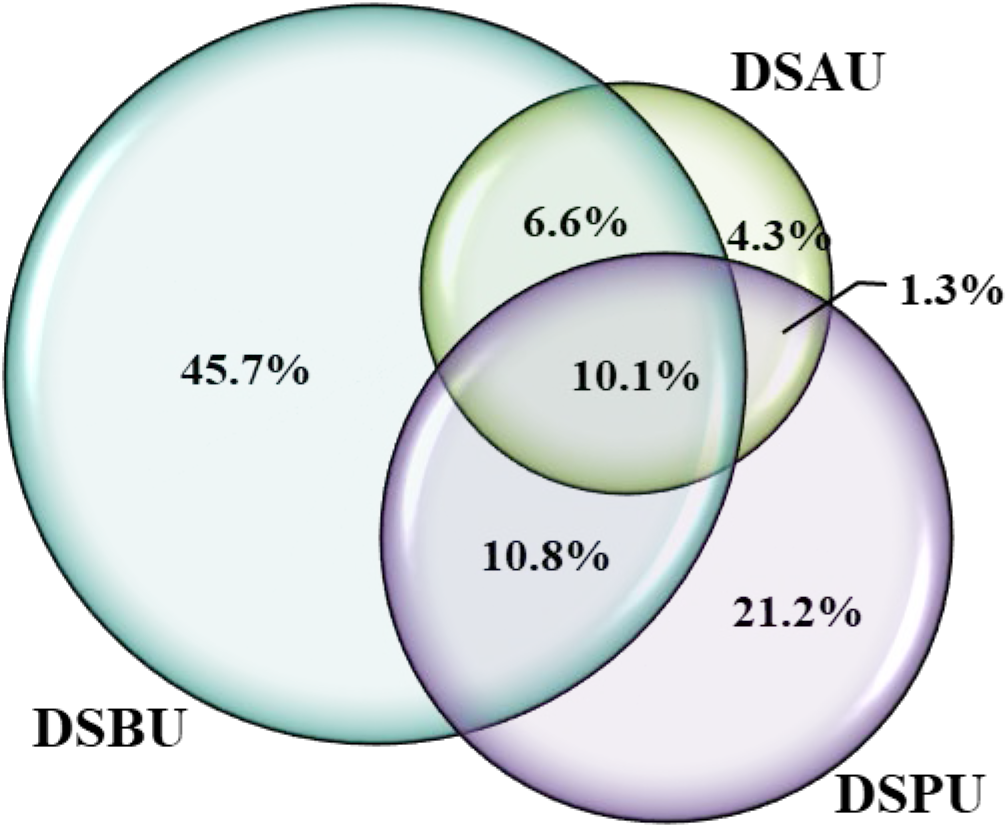
Overall overlap of BSA cross-links for DSBU, DSPU, and DSAU. All cross-linked products identified from three technical replicates with three cross-linkers using three concentrations are included.

The average Cα-Cα distances bridged by the three cross-linkers range between 18.9 Å for DSAU, 20.9 Å for DSPU, and 21.2 Å for DSBU (SI Figure S2), which is in perfect agreement with the different spacer lengths of the three cross-linkers (Scheme 1). For all three cross-linkers, the maximum Cα-Cα distance bridged is ~ 35 (40) Å, which harmonizes with our previous studies^3, 6, 21^. All unique cross-linking sites of BSA are summarized in SI Tables.

### Cross-links in E. coli Ribosome

For cross-linking experiments of the *E. coli* ribosome, we used exclusively DSBU as cross-linker. In total, 868 cross-linking sites were identified for the *E. coli* ribosome (Table 2, Figure 4A). The results with DSBU agree well with the data acquired recently in our group for the *E. coli* ribosome using the shorter DSAU linker^21^: 55% of unique cross-linking sites were found in both studies. The four replicates show an overall overlap of 34% with 295 unique cross-linking sites being identified in all four experiments (Figure 4B). All unique cross-linking sites in the *E. coli* ribosome are summarized in SI Tables.

**Figure 4:**
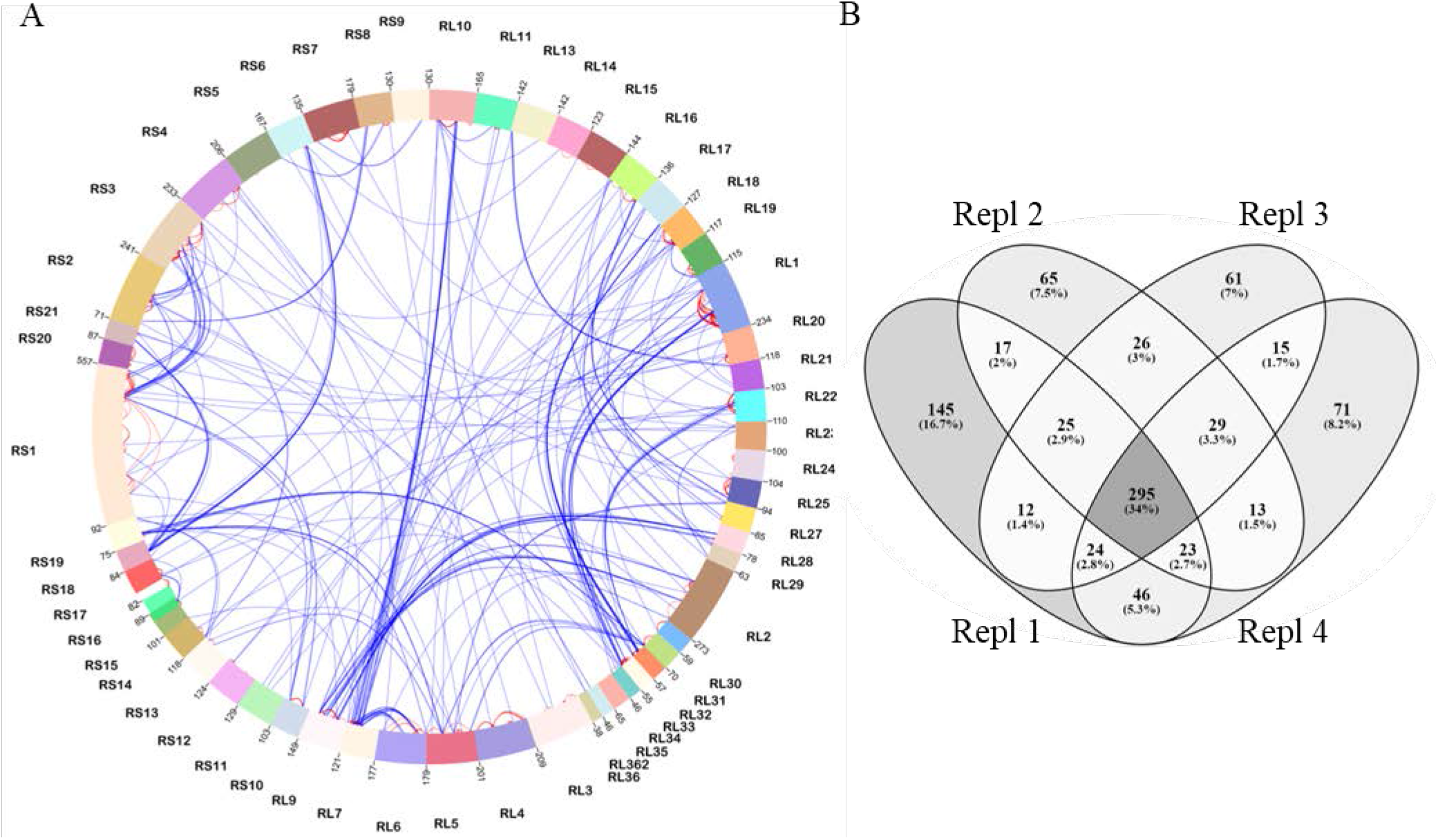
Cross-linking sites identified in *E. coli* ribosome. (A) Cross-linking sites identified in the *E. coli* ribosome are presented as Circular plot with xVis (https://xvis.genzentrum.lmu.de). Intermolecular cross-links between different ribosomal subunits are shown in blue, intramolecular cross-links within one ribosomal subunit are shown in red. RL: large subunit; RS: small subunit; (B) Venn diagram showing the overlap of cross-linking sites identified in four replicates.

**Table 2:** Unique cross-linking sites and cross-link spectral matches (CSMs) for *E. coli* ribosome. Numbers are given for four replicates (DSBU, 50-fold molar excess) as well as the combined numbers.

|  | **CSMs** | **Unique sites** |
| --- | --- | --- |
| **Repl 1** | 3407 | 587 |
| **Repl 2** | 2998 | 493 |
| **Repl 3** | 2984 | 488 |
| **Repl 4** | 3537 | 517 |
| **combined** | 12926 | 868 |

### Protein-Protein Interactions in Nuclear Lysates

For the XL-MS experiments, HEK293T cells were first incubated with a lysis buffer containing digitonin to keep the nuclear protein-protein interactions intact before the cross-linking reaction. Digitonin is a mild detergent to facilitate cell lysis via cell membrane permeabilization without denaturing proteins. After centrifugation, the cross-linking reaction was performed before breaking the nucleus membrane. Therefore, the nuclear lysates also contain proteins from other organelles like mitochondria, endoplasmatic reticulum, and Golgi. For searching cross-linked products of proteins in the HEK293T cell nuclear lysates, we did not consider any post-translational modifications, such as phosphorylation, glycosylation, acetylation or methylation, nor any truncations. For an in-depth analysis of the nuclear lysates, cross-linked samples were fractionated by SEC to enrich cross-linked peptides^14^ (Figure 5A). Fractions enriched in cross-linked peptides (B1 to B7) were directly subjected to LC/MS/MS without any further sample preparation. The numbers of total unique cross-linking sites identified in these fractions range between 51 (fraction B1) and 885 (fraction B3; Figure 5B) with an overlap between the three replicates of ~ 17% (Figure 5C). All unique cross-linking sites of nuclear lysate samples are summarized in SI Tables.

**Figure 5:**
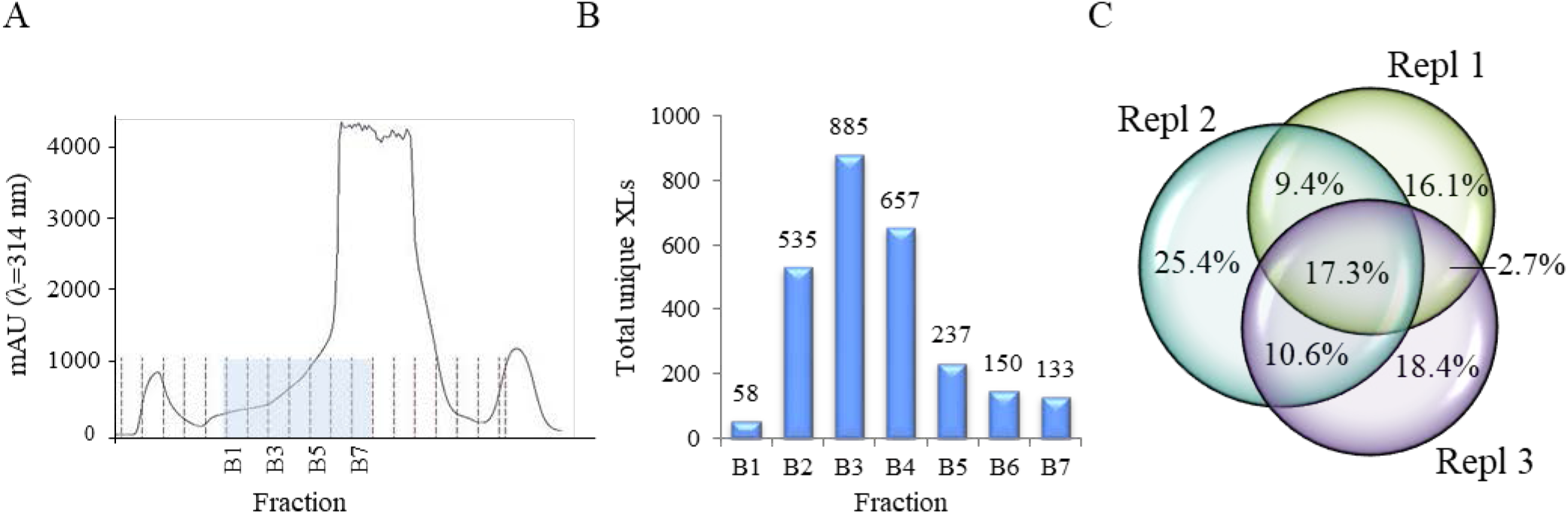
Cross-link analysis of HEK293T cell nuclear lysates. (A) SEC separation of one nuclear lysate sample; (B) unique cross-linking sites from three biological replicates contained in selected SEC fractions; (C) Venn diagram showing the overlap of cross-linking sites identified in three replicates.

In total, 1,623 unique cross-linking sites corresponding to 564 potential protein-protein interactions were identified in HEK293T cell nuclear lysates (Figure 6). In the following, some of these interactions will be highlighted.

**Figure 6:**
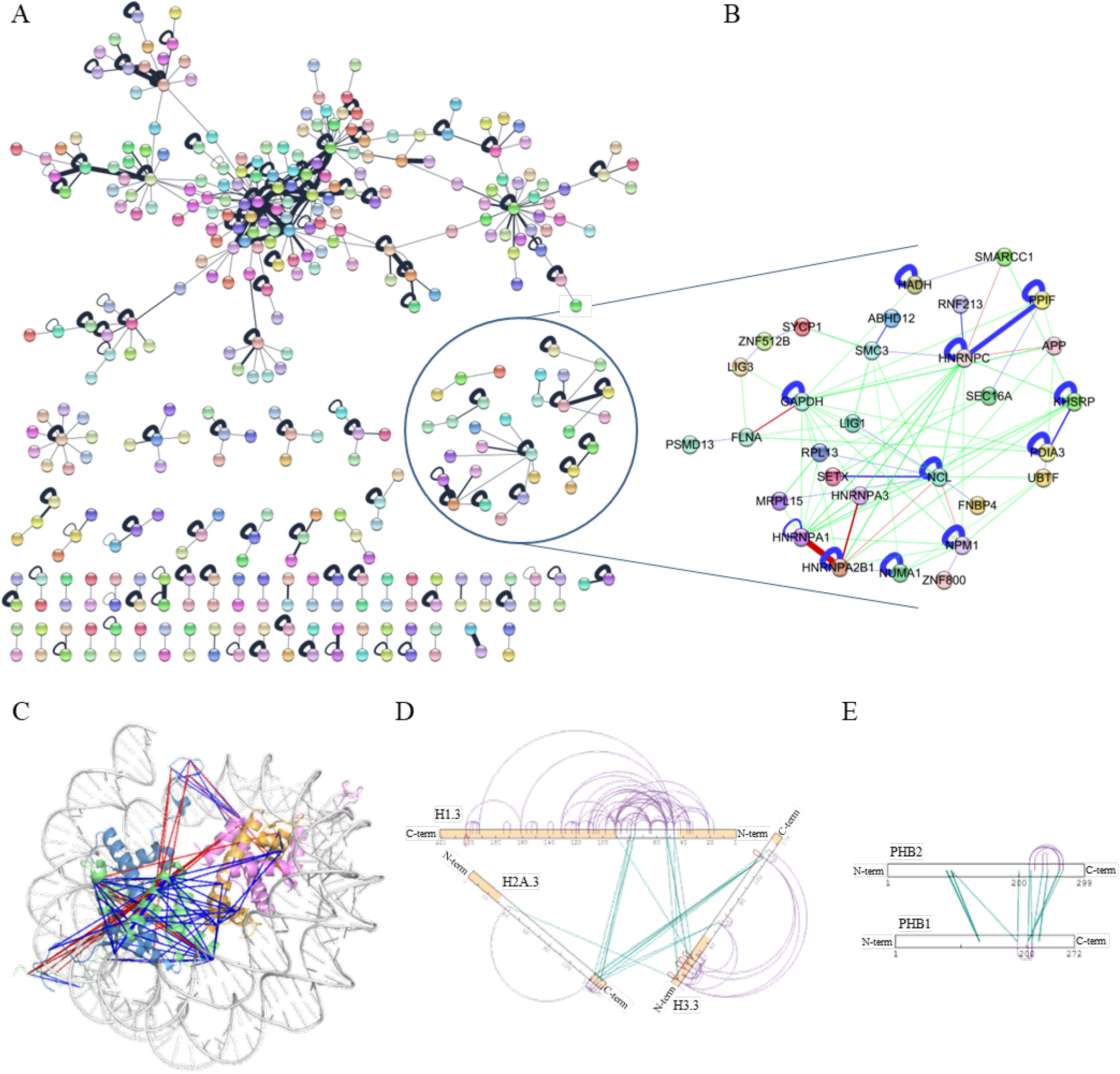
Protein-protein interaction network in HEK293T cell nuclear lysates. (A) Unique cross-linking sites and potential protein-protein interactions identified are schematically presented. The thickness of the lines corresponds to the number of cross-links; (B) zoom-in view on the protein cluster involved in nucleic acid processing; (C) unique cross-linking sites for the four histones of the nucleosome, H2A, H2B, H3, and H4 (pdb entry 1EQZ), cross-links are indicated in blue: ≤ 40 Å and in red: > 40 Å; (D) intra- and interprotein cross-linking data of H1.3, H2A type 3, and H3.3 as well as (E) intra- and interprotein cross-linking data of PHB1 and PHB2 are shown schematically with xiNET (http://crosslinkviewer.org); intramolecular cross-links are shown in pink, intermolecular cross-links in green; homotypic cross-links in red. Intrinsically disordered regions in the histones are presented in orange.

### Proteins of Nucleic Acid Processing

We identified five clusters with 30 protein involved in nucleic acid processing (Figure 6B). According to the STRING database (https://string-db.org) these proteins are all interconnected in a larger and intricate protein network. Our XL-MS data confirmed several known interactions and also detected a number of novel protein-protein interactions, involving nucleolin, hnRNP, probable helicase senataxin, nuclear mitotic apparatus protein 1, and nucleophosmin.

### Histone Proteins

Another cluster of nuclear protein-protein interactions was discovered for histone proteins. Here, we discovered 105 unique cross-linking sites for the four histones of the nucleosome, H2A, H2B, H3, and H4 (Figure 6C). In total, 86 cross-links were below the Cα-Cα limit below 40 Å, according to pdb structure 1EQZ for the nucleosome. The 19 cross-links that exceed the maximum Cα-Cα distance of 40 Å can be explained by the highly flexible, intrinsically disordered regions in the respective histones. Alternatively, these “overlength” cross-links could also originate from connecting another histone monomer in the complex as each of the four histones, H2A, H2B, H3, and H4, is present twice in the complex^25^. Our data reveal cross-links involving the intrinsically disordered regions in H2A and H3 that are not resolved in the X-ray structure (pdb 1EQZ)^26^. In another study however the complete structure of H3 is resolved^27^. The cross-links with the intrinsically disordered regions of H3 to other components of the nucleosome are longer than 40 Å, which is easily explained by the high flexibility of the regions involved. This underlines the strength of the XL-MS approach to gain 3D-structural information on intrinsically disordered regions. Histone H1 is present in a few X-ray and cryo-EM structures of the chromatosome (histone complex) [pdb entries 7K5X, 6L9Z, 5NL0], yet only with its globular domain, while the flexible part of H1 is missing in these structures. Again, our intraprotein cross-linking data of H1 (Figure 6D) cover the intrinsically disordered regions of H1. Cross-links identified in the histone isoform H1.3 clearly show an orientation of cross-linked H1.3 at the chromatosome where its globular domain interacts with the intrinsically disordered *C*- and *N*-termini of histones H2A and H3. Strikingly, no cross-links were found between the nucleosome and the long intrinsically disordered C-terminus of H1.3, confirming the orientation of H1.3 along the linker DNA as has been observed previously^28^.

### Prohibitin Complexes

Another protein-protein interaction discovered in our XL-MS studies was the complex between PHB1 and PHB2 (Figure 6E). PHB1-PHB2 interactions have recently been studied by XL-MS in mitochondria^29^. We identified this complex as it is also present in nucleus and we cannot completely exclude the presence of mitochondrial proteins in our preparation. Interestingly, the cross-links found between PHB1 and PHB2 in that previous study are in perfect agreement to our findings, connecting identical sites in both proteins. This underlines the power of our approach in identifying protein interaction sites in a highly complex environment.

## CONCLUSION

We show that combining XL-MS with IM-MS can be employed without sophisticated CCS selection methods for a rapid and thorough investigation of cross-linking mixtures, ranging from single proteins to highly complex protein mixtures. The benefits for analyzing biological samples are high as instrumental settings do not have to be changed for each protein system under investigation. Therefore, we envision our strategy to play an important role in developing routine strategies for structural proteomics studies.

## Abbreviations

ACN: Acetonitrile
BSA: Bovine serum albumin
Caps: Collisional cross-section assisted precursor selection
CCS: Collisional cross section
CSM: Cross-link spectral match
DSAU: Disuccinimidyl diacetic urea
DSBU: Disuccinimidyl dibutyric urea
DSPU: Disuccinimidyl dipropionic urea
DTT: Dithiothreitol
EM: Electron microscopy
FDR: False discovery rate
HEPES: 2-[4-(2-Hydroxyethyl)-piperazine-1-yl]ethanesulfonic acid
IM: Ion mobility
IMAC: Immobilized metal ion chromatography
LC: Liquid chromatography
MS: Mass spectrometry
MS/MS: Tandem mass spectrometry
NHS: *N*-Hydroxysuccinimide
PASEF: Parallel accumulation serial fragmentation
PBS: Phosphate buffered saline
RT: Room temperature
SAXS: Small-angle X-ray scattering
SDS: Sodium dodecyl sulfate
SEC: Size exclusion chromatography
TIMS: Trapped ion mobility spectrometry
TOF: Time-of-flight
XL: Cross-linking
XL-MS: Cross-linking/mass spectrometry

## ASSOCIATED CONTENT

### Supporting Information

The Supporting Information contains the Venn diagrams for each replicate of BSA cross-linking (Figure S1) and the distribution of Cα-Cα distances in BSA for the three cross-linkers (Figure S2). Tables of all unique cross-links in BSA, *E. coli* ribosome and HEK293T cell nuclear lysates are presented in one xlsx file.

## ACKNOWLEDGEMENTS

A.S. acknowledges financial support by the DFG (INST 271/404-1 FUGG, INST 271/405-1 FUGG, and RTG 2467, project number 391498659 “Intrinsically Disordered Proteins – Molecular Principles, Cellular Functions, and Diseases”), the region of Saxony-Anhalt, and the Martin Luther University Halle-Wittenberg (Center for Structural Mass Spectrometry). The authors are indebted to Dr. Marcel Köhn for cell culture and preparation of nuclear lysates. Dr. Markus Lubeck, Dr. Gary Kruppa, and Dr. Florian Busch (all Bruker Daltonics) are acknowledged for helpful discussions. The authors would like to thank Department 4 of the Martin Luther University Halle-Wittenberg for the reconstruction of the former “Tierstall”.

## Author Contributions

All authors have given approval to the final version of the manuscript.

---

**Scheme 1:**
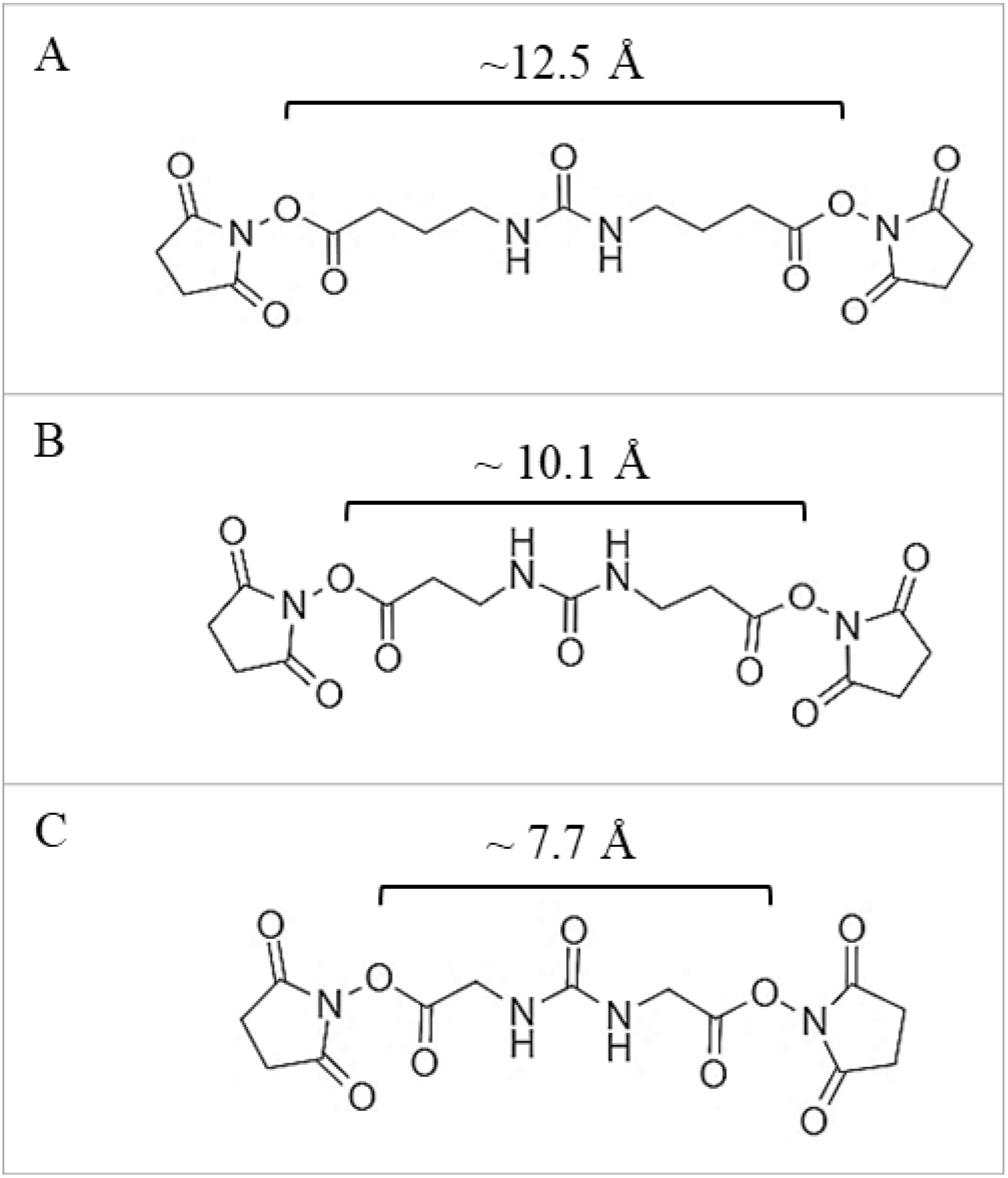
Structures and spacer lengths of urea cross-linkers. All cross-linkers contain *N*-hydroxysuccinimide (NHS) ester as reactive groups. The carbon spacer arm contains four, three, and two C-atoms on each side of the cleavable urea group for (A) DSBU, (B) DSPU, and (C) DSAU.

## References

1. Schmidt, C.; Urlaub, H., Combining cryo-electron microscopy (cryo-EM) and cross-linking mass spectrometry (CX-MS) for structural elucidation of large protein assemblies. Current opinion in structural biology 2017, 46, 157–168.

2. O’Reilly, F. J.; Rappsilber, J., Cross-linking mass spectrometry: methods and applications in structural, molecular and systems biology. Nature structural & molecular biology 2018, 25 (11), 1000–1008.

3. Iacobucci, C.; Piotrowski, C.; Aebersold, R.; Amaral, B. C.; Andrews, P.; Bernfur, K.; Borchers, C.; Brodie, N. I.; Bruce, J. E.; Cao, Y.; Chaignepain, S.; Chavez, J. D.; Claverol, S.; Cox, J.; Davis, T.; Degliesposti, G.; Dong, M. Q.; Edinger, N.; Emanuelsson, C.; Gay, M.; Gotze, M.; Gomes-Neto, F.; Gozzo, F. C.; Gutierrez, C.; Haupt, C.; Heck, A. J. R.; Herzog, F.; Huang, L.; Hoopmann, M. R.; Kalisman, N.; Klykov, O.; Kukacka, Z.; Liu, F.; MacCoss, M. J.; Mechtler, K.; Mesika, R.; Moritz, R. L.; Nagaraj, N.; Nesati, V.; Neves-Ferreira, A. G. C.; Ninnis, R.; Novak, P.; O’Reilly, F. J.; Pelzing, M.; Petrotchenko, E.; Piersimoni, L.; Plasencia, M.; Pukala, T.; Rand, K. D.; Rappsilber, J.; Reichmann, D.; Sailer, C.; Sarnowski, C. P.; Scheltema, R. A.; Schmidt, C.; Schriemer, D. C.; Shi, Y.; Skehel, J. M.; Slavin, M.; Sobott, F.; Solis-Mezarino, V.; Stephanowitz, H.; Stengel, F.; Stieger, C. E.; Trabjerg, E.; Trnka, M.; Vilaseca, M.; Viner, R.; Xiang, Y.; Yilmaz, S.; Zelter, A.; Ziemianowicz, D.; Leitner, A.; Sinz, A., First Community-Wide, Comparative Cross-Linking Mass Spectrometry Study. Analytical chemistry 2019, 91 (11), 6953–6961.

4. Klykov, O.; Steigenberger, B.; Pektas, S.; Fasci, D.; Heck, A. J. R.; Scheltema, R. A., Efficient and robust proteome-wide approaches for cross-linking mass spectrometry. Nature protocols 2018, 13 (12), 2964–2990.

5. Leitner, A.; Faini, M.; Stengel, F.; Aebersold, R., Crosslinking and Mass Spectrometry: An Integrated Technology to Understand the Structure and Function of Molecular Machines. Trends in biochemical sciences 2016, 41 (1), 20–32.

6. Iacobucci, C.; Gotze, M.; Ihling, C. H.; Piotrowski, C.; Arlt, C.; Schafer, M.; Hage, C.; Schmidt, R.; Sinz, A., A cross-linking/mass spectrometry workflow based on MS-cleavable cross-linkers and the MeroX software for studying protein structures and protein-protein interactions. Nature protocols 2018, 13 (12), 2864–2889.

7. Yu, C.; Huang, L., Cross-Linking Mass Spectrometry: An Emerging Technology for Interactomics and Structural Biology. Analytical chemistry 2018, 90 (1), 144–165.

8. Chavez, J. D.; Bruce, J. E., Chemical cross-linking with mass spectrometry: a tool for systems structural biology. Current opinion in chemical biology 2019, 48, 8–18.

9. Petrotchenko, E. V.; Borchers, C. H., Crosslinking combined with mass spectrometry for structural proteomics. Mass spectrometry reviews 2010, 29 (6), 862–76.

10. Yang, B.; Wu, Y. J.; Zhu, M.; Fan, S. B.; Lin, J.; Zhang, K.; Li, S.; Chi, H.; Li, Y. X.; Chen, H. F.; Luo, S. K.; Ding, Y. H.; Wang, L. H.; Hao, Z.; Xiu, L. Y.; Chen, S.; Ye, K.; He, S. M.; Dong, M. Q., Identification of cross-linked peptides from complex samples. Nature methods 2012, 9 (9), 904–6.

11. Chavez, J. D.; Mohr, J. P.; Mathay, M.; Zhong, X.; Keller, A.; Bruce, J. E., Systems structural biology measurements by in vivo cross-linking with mass spectrometry. Nature protocols 2019, 14 (8), 2318–2343.

12. O’Reilly, F. J.; Xue, L.; Graziadei, A.; Sinn, L.; Lenz, S.; Tegunov, D.; Blotz, C.; Singh, N.; Hagen, W. J. H.; Cramer, P.; Stulke, J.; Mahamid, J.; Rappsilber, J., In-cell architecture of an actively transcribing-translating expressome. Science 2020, 369 (6503), 554–557.

13. Liu, F.; Lossl, P.; Rabbitts, B. M.; Balaban, R. S.; Heck, A. J. R., The interactome of intact mitochondria by cross-linking mass spectrometry provides evidence for coexisting respiratory supercomplexes. Molecular & cellular proteomics: MCP 2018, 17 (2), 216–232.

14. Gotze, M.; Iacobucci, C.; Ihling, C. H.; Sinz, A., A Simple Cross-Linking/Mass Spectrometry Workflow for Studying System-wide Protein Interactions. Analytical chemistry 2019, 91 (15), 10236–10244.

15. Silveira, J. A.; Ridgeway, M. E.; Laukien, F. H.; Mann, M.; Park, M. A., Parallel accumulation for 100% duty cycle trapped ion mobility-mass spectrometry. International Journal of Mass Spectrometry 2017, 413, 168–175.

16. Meier, F.; Beck, S.; Grassl, N.; Lubeck, M.; Park, M. A.; Raether, O.; Mann, M., Parallel Accumulation-Serial Fragmentation (PASEF): Multiplying Sequencing Speed and Sensitivity by Synchronized Scans in a Trapped Ion Mobility Device. Journal of proteome research 2015, 14 (12), 5378–87.

17. Meier, F.; Brunner, A. D.; Koch, S.; Koch, H.; Lubeck, M.; Krause, M.; Goedecke, N.; Decker, J.; Kosinski, T.; Park, M. A.; Bache, N.; Hoerning, O.; Cox, J.; Rather, O.; Mann, M., Online Parallel Accumulation-Serial Fragmentation (PASEF) with a Novel Trapped Ion Mobility Mass Spectrometer. Molecular & cellular proteomics: MCP 2018, 17 (12), 2534–2545.

18. Yu, F.; Haynes, S. E.; Teo, G. C.; Avtonomov, D. M.; Polasky, D. A.; Nesvizhskii, A. I., Fast Quantitative Analysis of timsTOF PASEF Data with MSFragger and IonQuant. Molecular & cellular proteomics: MCP 2020, 19 (9), 1575–1585.

19. Steigenberger, B.; van den Toorn, H. W. P.; Bijl, E.; Greisch, J. F.; Rather, O.; Lubeck, M.; Pieters, R. J.; Heck, A. J. R.; Scheltema, R. A., Benefits of Collisional Cross Section Assisted Precursor Selection (caps-PASEF) for Cross-linking Mass Spectrometry. Molecular & cellular proteomics: MCP 2020, 19 (10), 1677–1687.

20. Muller, M. Q.; Dreiocker, F.; Ihling, C. H.; Schafer, M.; Sinz, A., Cleavable cross-linker for protein structure analysis: reliable identification of cross-linking products by tandem MS. Analytical chemistry 2010, 82 (16), 6958–68.

21. Tuting, C.; Iacobucci, C.; Ihling, C. H.; Kastritis, P. L.; Sinz, A., Structural analysis of 70S ribosomes by cross-linking/mass spectrometry reveals conformational plasticity. Scientific reports 2020, 10 (1), 12618.

22. Grimm, M.; Zimniak, T.; Kahraman, A.; Herzog, F., xVis: a web server for the schematic visualization and interpretation of crosslink-derived spatial restraints. Nucleic acids research 2015, 43 (W1), W362–9.

23. Combe, C. W.; Fischer, L.; Rappsilber, J., xiNET: cross-link network maps with residue resolution. Molecular & cellular proteomics: MCP 2015, 14 (4), 1137–47.

24. Shannon, P.; Markiel, A.; Ozier, O.; Baliga, N. S.; Wang, J. T.; Ramage, D.; Amin, N.; Schwikowski, B.; Ideker, T., Cytoscape: a software environment for integrated models of biomolecular interaction networks. Genome research 2003, 13 (11), 2498–504.

25. Kornberg, R. D., Chromatin Structure: A Repeating Unit of Histones and DNA. Science 1974, (184).

26. Harp, J. M.; Hanson, B. L.; Timm, D. E.; Bunick, G. J., Asymmetries in the nucleosome core particle at 2.5 A resolution. Acta crystallographica. Section D, Biological crystallography 2000, 56 (Pt 12), 1513–34.

27. Davey, C. A.; Sargent, D. F.; Luger, K.; Maeder, A. W.; Richmond, T. J., Solvent Mediated Interactions in the Structure of the Nucleosome Core Particle at 1.9Å Resolution. Journal of Molecular Biology 2002, 319 (5), 1097–1113.

28. Bednar, J.; Garcia-Saez, I.; Boopathi, R.; Cutter, A. R.; Papai, G.; Reymer, A.; Syed, S. H.; Lone, I. N.; Tonchev, O.; Crucifix, C.; Menoni, H.; Papin, C.; Skoufias, D. A.; Kurumizaka, H.; Lavery, R.; Hamiche, A.; Hayes, J. J.; Schultz, P.; Angelov, D.; Petosa, C.; Dimitrov, S., Structure and Dynamics of a 197 bp Nucleosome in Complex with Linker Histone H1. Molecular cell 2017, 66 (3), 384–397 e8.

29. Yoshinaka, T.; Kosako, H.; Yoshizumi, T.; Furukawa, R.; Hirano, Y.; Kuge, O.; Tamada, T.; Koshiba, T., Structural Basis of Mitochondrial Scaffolds by Prohibitin Complexes: Insight into a Role of the Coiled-Coil Region. iScience 2019, 19, 1065–1078.

