## Supporting Information, Figures S1 and S2 for "Cross-linking/Mass Spectrometry Combined with Ion Mobility on a timsTOF Pro Instrument for Structural Proteomics"

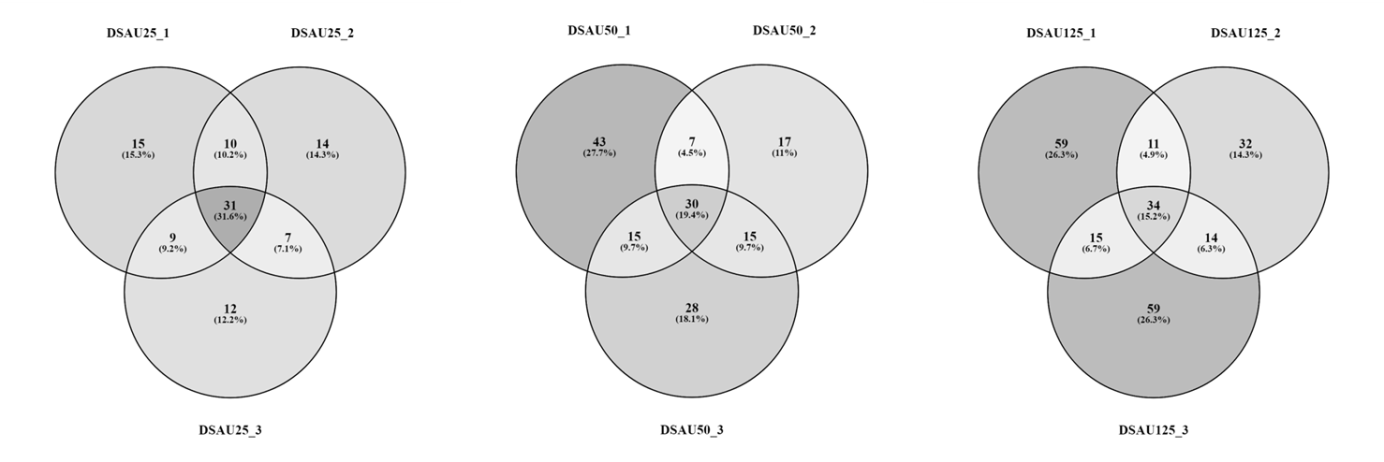

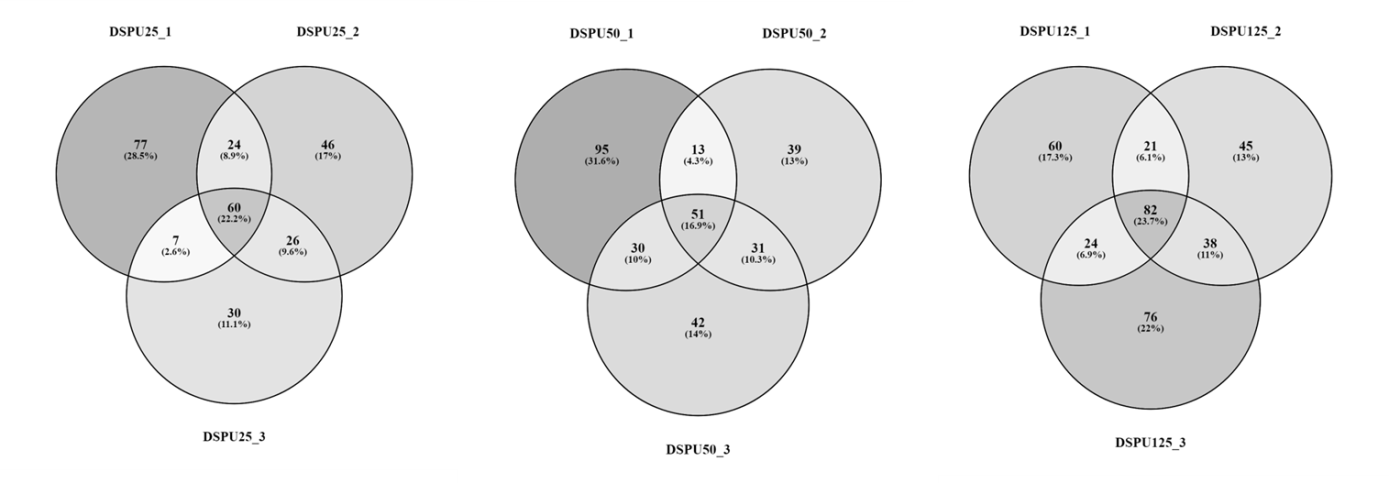

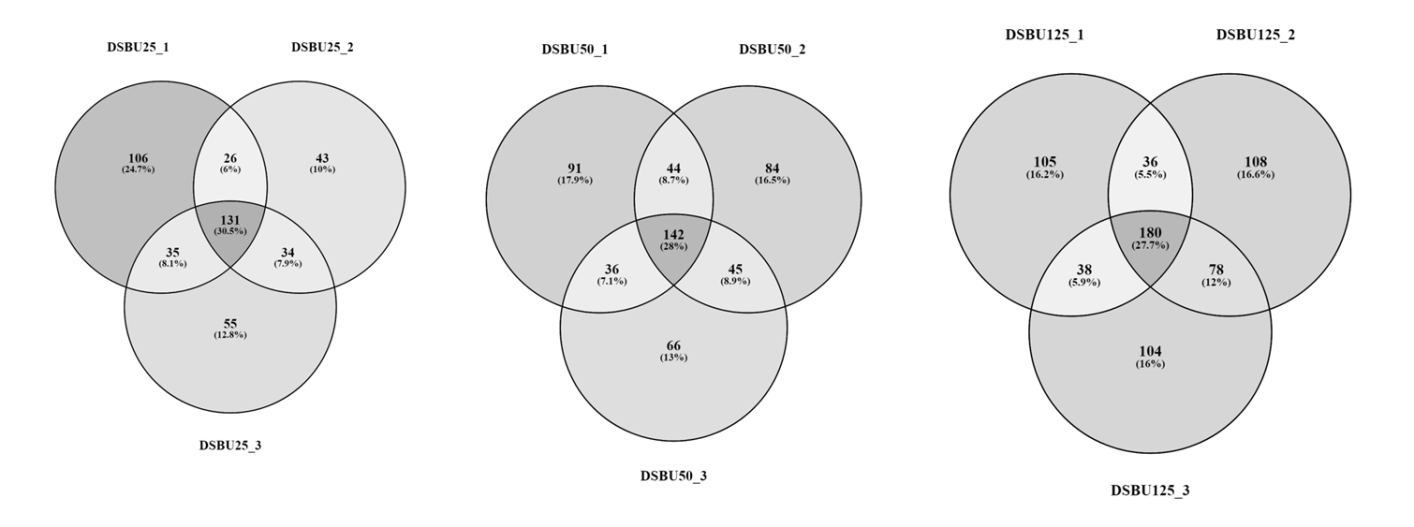

**Figure S1:**	**Venn diagrams for each BSA cross-linking replicate**. Cross-linker names (DSAU, DSPU, DSBU), molar excess of cross-linker (25, 50, 125), and replicate numbers (1-3) are given.

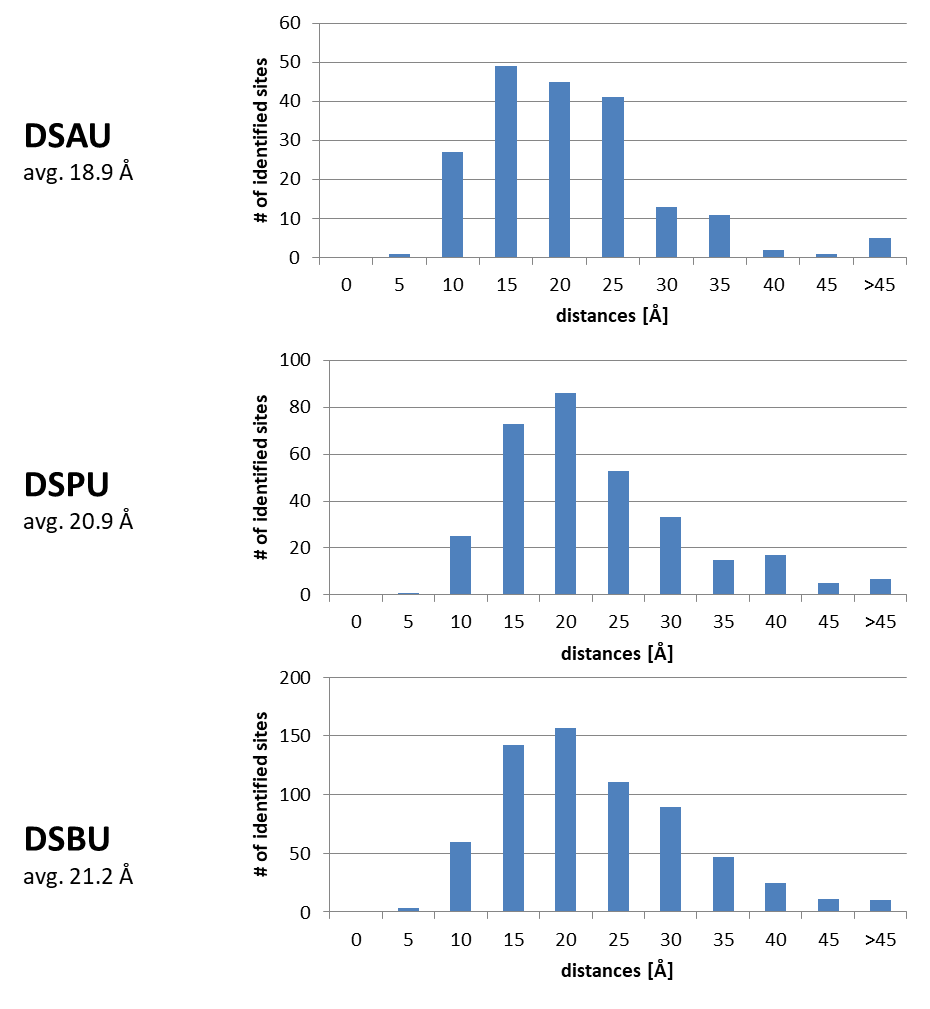

**Figure S2**:	**Distribution of Cα-Cα distances in BSA for the three cross-linkers used in this work.** For spacer lengths of the cross-linkers DSBU, DSPU, and DSAU please see Scheme 1.
